# Convergence and function of distinct corticocortical and transthalamic synapses across sensory processing hierarchies

**DOI:** 10.64898/2026.09.08.750213

**Authors:** Mckenzie Haller, Garrett T. Neske

## Abstract

Communication among cortical regions underlies fundamental functions of the brain. Thus, it is essential to understand the synaptic mechanisms by which distinct cortical regions become functionally connected. While direct corticocortical synaptic connections between cortical regions have traditionally been considered the primary mode of cortex-wide interaction, these regions can also interact indirectly through transthalamic pathways that engage higher-order regions of the thalamus. Yet, the mechanisms by which these two distinct inter-cortical synaptic pathways exert their influence on their target cortical neurons are largely unknown. A major obstacle to the targeted synaptic analysis of these pathways has been the lack of an appropriate method to genetically label them, especially in the same preparation. Here, using a combination of viral vector methods for anterograde transsynaptic and retrograde labeling along with optogenetic activation of long-range synaptic terminals in *ex vivo* slices of mouse visual cortical areas, we uncover a number of previously unknown features of corticocortical pathways and their transthalamic counterparts. First, corticocortical and transthalamic pathways originating from the same cortical source are largely monosynaptically integrated by target cortical neurons, though with an integration frequency that depends on the hierarchical level of the sensory processing stream. Second, feedforward transthalamic and corticocortical synapses are dynamically distinct and activate responses with distinct postsynaptic receptor composition, whereas these same pathways in the feedback direction elicit synaptic responses that are largely comparable. Overall, these results provide a novel synaptic-level framework for understanding how cortical and thalamic axon pathways shape the flow of information across interareal cortical networks.

## INTRODUCTION

Synaptic interactions among the many regions of the neocortex are foundational for normal operations of the brain, including sensory perception, motor control, and cognition. Cortex-wide synaptic signaling sculpts incoming sensory data to build complex, coherent percepts (Orban, 2008; Riesenhuber and Poggio, 1999). Furthermore, dynamic alterations of this signaling underlie context-dependent sensory processing, shifting the moment-to-moment functional connectivity of local and long-range cortical circuits to support cognition (Cardin, 2019; Nakajima and Halassa, 2017; Haider and McCormick, 2009). A significant component of cortical synaptic signaling is hierarchical in nature, with synaptic terminals originating from lower-order cortices carrying information about more basic sensory features to be combined by higher-order cortical neurons to generate novel feature selectivity (El-Shamayleh *et al*., 2013; Movshon and Newsome, 1996). Simultaneously, these higher-order cortical neurons provide various forms of contextual modulation in the form of feedback afferents to lower-order cortical neurons (Keller *et al*., 2020; Gilbert and Li, 2013; Bastos *et al*., 2012). Given the prevalence of hierarchical interactions within the cortex, it is essential to determine their synaptic mechanisms (short-term dynamics of specific synaptic terminals, postsynaptic receptors engaged, dendritic integration principles, etc.).

Traditional models of cortex-wide hierarchical processing consider the direct synaptic connections between cortical neurons (corticocortical connections) to be paramount. Yet, the role of the thalamus in dynamic functional cortical connectivity is becoming increasingly acknowledged, particularly the contributions of *transthalamic pathways* – indirect connections between cortical regions that engage higher-order regions of the thalamus as an intermediary (Sherman and Usrey, 2024). In contrast to first-order thalamus, which receives its primary driving synaptic input from more peripheral structures, higher-order thalamus receives such input from the cortex. Earlier anatomical evidence suggested the principle that any two cortical regions connected directly via corticocortical connections are also connected via transthalamic pathways, such as by relay cells in the visual pulvinar nucleus (Shipp 2003; Guillery, 1995). Recent evidence in a variety of preparations indicates that transthalamic pathways, alongside their corticocortical counterparts, convey crucial sensory and contextual information across large-scale cortical networks (Neske and Cardin, 2025; McKinnon *et al*., 2025; Mo *et al*., 2024; Blot *et al*. 2021). Yet, we still lack many key synaptic- and circuit-level details regarding the mechanisms of corticocortical and transthalamic transmission in hierarchically defined cortical regions. Indeed, unlike for first-order thalamocortical systems and their associated primary sensory cortices (Castro-Alamancos, 2004; Amitai, 2001), fundamental knowledge of the synaptic efficacy of the cortical and thalamic pathways interconnecting different cortical regions has lagged. Furthermore, despite the hypothesized possibility that the coincidence of corticocortical and transthalamic signals in the cortex might be optimal for cortical functional connectivity changes (Sherman and Guillery, 2013 & 2011), we lack any data on the mechanisms of synaptic integration of these two types of signals.

Investigations of the comparative synaptic impact of corticocortical and transthalamic pathways on their cortical target neurons require methods for selective activation of these pathways. Additionally, determination of the mechanisms of corticocortical-transthalamic integration within the cortex necessitates a procedure by which these pathways can be independently stimulated within one experimental preparation. Here, harnessing a combination of viral-based intersectional genetic techniques and optogenetics, we fulfill both of these requirements to uncover the synaptic properties of corticocortical and partnering transthalamic pathways mediated by the pulvinar nucleus in both feedforward and feedback directions at multiple hierarchical processing levels of the mouse visual system. Well-studied with respect to the functional relationships among its various cortical regions (Glickfeld and Olsen, 2017), the mouse visual system provides an ideal platform to explore the relationships between corticocortical and transthalamic interactions at distinct processing stages. We show that corticocortical and transthalamic pathways originating from the same source cortex are monosynaptically integrated by neurons in the target cortex, but with integration prevalence and synaptic properties that differ depending on the direction of information flow (feedforward vs. feedback) and level of the processing hierarchy. Our results provide a novel synaptic- and circuit-level framework for integrating thalamic contributions into models of dynamic functional cortical connectivity.

## RESULTS

### Genetic labeling of higher-order thalamic relay cells participating in specific transthalamic pathways

While cortical regions are well-known to exhibit interconnectivity mediated by long-range, monosynaptic linkages between cortical neurons (Harris and Shepherd, 2015; Felleman and Van Essen, 1991), higher-order thalamic regions such as the visual pulvinar nucleus provide additional conduits for cortex-wide information flow (Sherman and Guillery, 2013). Indeed, it is conceivable that every direct corticocortical connection is mirrored by a corresponding disynaptic *transthalamic* connection consisting of a corticothalamic synapse made in a higher-order thalamic nucleus, followed by a thalamocortical synapse made in another cortical region (Shipp, 2003). A major obstacle to studying the comparative synaptic impact and functional roles of a given transthalamic pathway that shares the same cortical source as a given corticocortical pathway has been the lack of a method to label such disynaptically defined thalamic relay cells. Here, for the first time, we describe such a method by combining anterograde transsynaptic viral tracing with intersectional genetics. We implement this novel technique in several hierarchically defined cortical areas of the mouse visual system, labeling pulvinar thalamic relay cells based upon their directional (i.e. feedforward or feedback) connectivity with these cortical areas. At the lowest hierarchical level, we label the feedforward and feedback transthalamic connections between primary visual cortex (V1) and the lateromedial visual cortex (LM) – considered the secondary visual cortex, V2, of the mouse (D’Souza *et al*., 2022; Wang and Burkhalter, 2007) (Fig. 1A). At the highest hierarchical level, we label the feedforward and feedback transthalamic connections between the posteromedial visual cortex (PM) and the anterior cingulate cortex (ACC) (Fig. 1A). We used high-titer AAV1-Cre injections in a source cortical region (e.g. V1) as an anterograde transsynaptic tracer to label the neurons postsynaptic to this source region with Cre recombinase (e.g. V1→pulvinar neurons in the pulvinar). As previously demonstrated in other systems (Zingg *et al*. 2020 & 2017), we find that pulvinar neurons (labeled with Cre after an AAV1-Cre injection in V1) respond monosynaptically to stimulation of the AAV1-Cre-labeled axons from V1 (Fig. 1B). Thus, with high-titer AAV1-Cre injection within a specific cortical region, pulvinar neurons receiving corticothalamic monosynaptic inputs from this region are labeled with Cre. To complete a defined transthalamic pathway, such Cre-labeled pulvinar neurons also need to be labeled based upon the cortical region to which they project. To fulfill this criterion, we also perform an injection of the retrogradely labeling CAV2-FLEx^loxP^-Flp virus in the target cortical region. Neurons whose axon terminals take up this virus will then express Cre-dependent Flp recombinase, such that, if these neurons also express Cre, they will thereby express Flp. Thus, by this method, pulvinar cell bodies that express Flp disynaptically connect two cortical regions in specified directions, beginning at the corticothalamic source where the AAV1-Cre injection was made and ending at the thalamocortical target where the CAV2-FLEx^loxP^-Flp injection was made (Fig. 1C).

**Figure 1.**
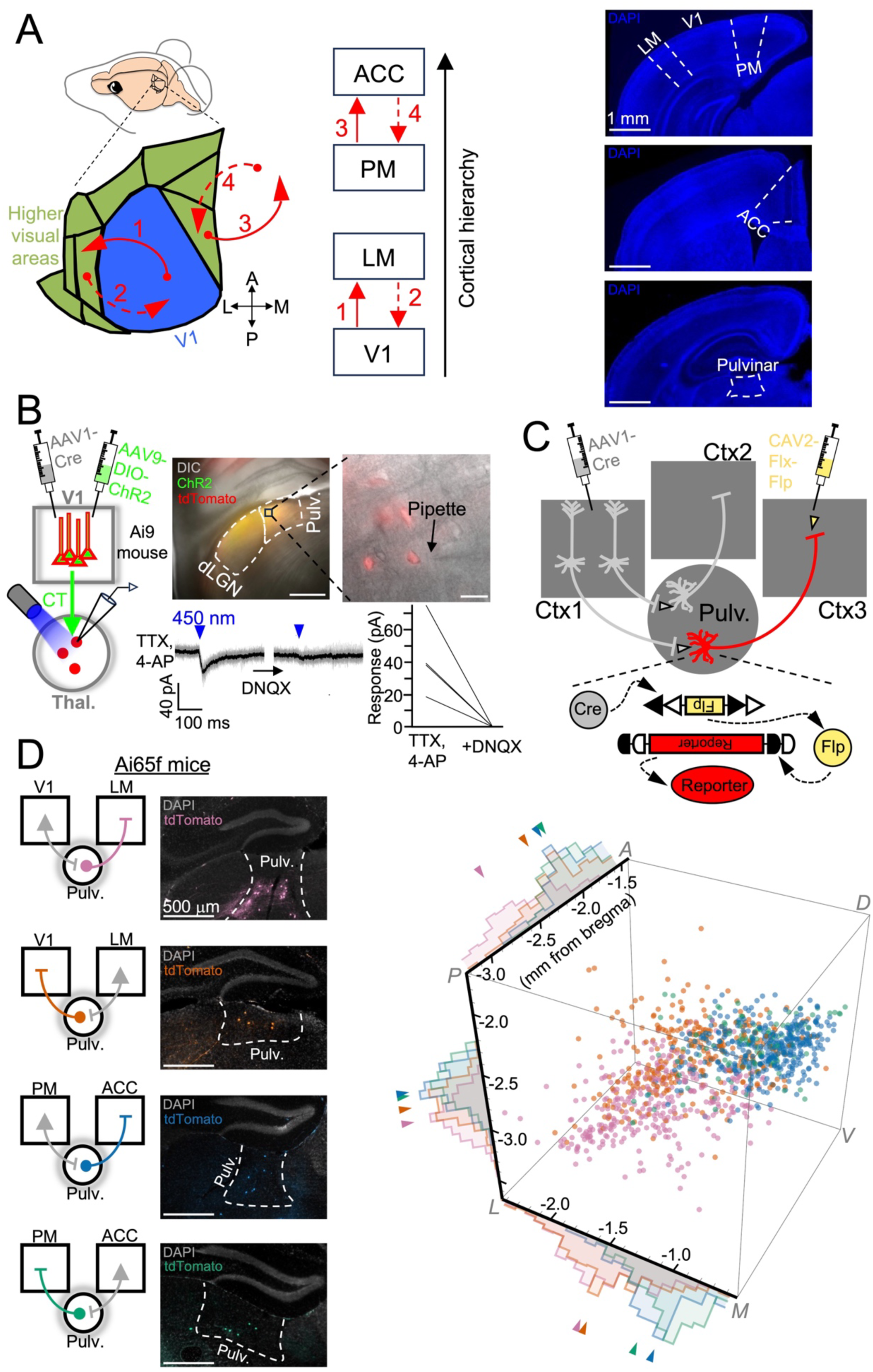
Genetic labeling of hierarchically defined visual transthalamic pathways. **(A)** (*Left*) Schematic of mouse visual cortical areas and their feedforward and feedback connections. V1 = primary visual cortex, LM = lateromedial visual cortex, PM = posteromedial visual cortex, ACC = anterior cingulate cortex. ACC is located within the longitudinal fissure of the mouse brain, relatively far from the occipital visual areas. (*Right*) Coronal histological sections demarcating the regions relevant to this study. **(B)** (*Left*) Schematic for optogenetic verification of anterograde transsynaptic infection of thalamic neurons by V1 cortical MV1-Cre injection. In the Ai9 (Cre-dependent tdTomato reporter) mouse, neurons in thalamus receiving monosynaptic corticothalamic input from V1 fluoresce with tdTomato. (*Right*) Image of a live slice demonstrating overlap of presynaptic, ere-positive corticothalamic terminals labeled with ChR2 and postsynaptic, ere-positive thalamic neurons labeled with tdTomato. Under high-magnification, individual tdTomato-positive postsynaptic pulvinar neurons can be targeted for electrophysiological recording. Low-mag scale bar = 500 µm, high-mag scale bar = 20 µm. Electrophysiological data underneath images demonstrate functional monosynaptic glutamatergic responses in *n=4* tdTomato-positive pulvinar neurons. **(C)** Schematic of viral vector injections combined with intersectional genetics for labeling pulvinar neurons in specific transthalamic pathways. **(D)** (*Left*) Coronal histological sections from Ai65f (Flp-dependent tdTomato reporter) mice injected with MV1-Cre and CAV2-Flx-Flp in the appropriate source and target cortical areas to label pulvinar neurons mediating specific transthalamic pathways. (*Right*) Population data plotting the stereotaxic coordinates of fluorescently-labeled pulvinar neurons mediating specific transthalamic pathways (V1→pulv.→LM: *N=4* mice, *n*=338 neurons; LM pulv.→V1: *N=5* mice, *n=299* neurons; PM→pulv.→ACC: *N* =4 mice, *n=360* neurons; ACC→pulv.→PM: *N=5* mice, *n*=115 neurons). Arrows for each anatomical axis indicate median markers. The following statistical comparisons of position along each anatomical axis were significant (Kruskal-Wallis test followed by Dunn’s post-hoc test with p-values corrected for multiple comparisons using the Benjamini-Hochberg procedure). *Anterior-posterior axis:* V1→pulv.→LM vs. LM→pulv.→V1, p≈0; V1→pulv.→LM vs. PM→pulv.→ACC, p≈0; V1→pulv.→LM vs. ACC→pulv.→PM, p≈0 LM→pulv.→V1 vs. ACC→pulv.→PM, p=1x10^-6^; LM→pulv.→V1 vs. PM→pulv.→ACC, p=7x10^-6^. *Medial-lateral axis:* V1→pulv.→LM vs. PM→pulv.→ACC, p≈0; V1→pulv.→LM vs. ACC→pulv.→PM, p≈0; LM→pulv.→V1 vs. ACC→pulv.→PM, p≈0; LM→pulv.→V1 vs. PM→pulv.→ACC, p≈0; PM→pulv.→ACC vs. ACC→pulv.→PM, p=0.0005. *Dorsal-ventral axis:* V1→pulv.→LM vs. LM→pulv.→V1, p≈0; V1→pulv.→LM vs. PM→pulv.→ACC, p≈0; V1→pulv.→LM vs. ACC→pulv.→PM, p≈0; LM→pulv.→V1 vs. ACC→pulv.→PM, p≈0; LM→pulv.→V1 vs. PM→pulv.→ACC, p=6x10^-7^.

Using this new technique, we uncovered the spatial organization of four hierarchically distinct transthalamic pathways within the mouse pulvinar: the feedforward and feedback connections between V1 and LM (V1→pulvinar→LM and LM→pulvinar→V1, respectively) and the feedforward and feedback connections between PM and ACC (PM→pulvinar→ACC and ACC→pulvinar→PM, respectively). We performed the relevant AAV1-Cre and CAV2-FLEx^loxP^-Flp injections within V1, LM, PM, or ACC of Ai65f (Flp-dependent tdTomato reporter) mice (Fig. 1D). Counts of tdTomato-positive cells in stereotaxic coordinates (see Materials and Methods) revealed several key features of the anatomical arrangement of these distinct transthalamic pathways within the pulvinar (Fig. 1D). First, in the medial-lateral axis, the pulvinar neurons mediating the feedforward and feedback transthalamic pathways between V1 and LM were largely clustered in the lateral aspect, whereas those mediating the pathways between PM and ACC were clustered in the medial aspect. Second, in the anterior-posterior axis, the transthalamic pathway at the lowest hierarchical level (V1→pulvinar→LM pathway) engaged the most posterior aspect of the pulvinar, while with increasing hierarchical level (LM→pulvinar→V1 followed by PM→pulvinar→ACC followed by ACC→pulvinar→PM), transthalamic pathways become progressively oriented in the anterior aspect. Finally, in the dorsal-ventral axis, pulvinar neurons belonging to the V1→pulvinar→LM pathway were located most ventrally, followed by the LM→pulvinar→V1 pathway, and with the pulvinar neurons linking PM and ACC located most dorsally.

### Laminar segregation of synaptic inputs from hierarchically defined corticocortical and transthalamic pathways

Having determined the spatial organization of the V1→pulvinar→LM, LM→pulvinar→V1, PM→pulvinar→ACC, and ACC→pulvinar→PM transthalamic pathways within the mouse pulvinar nucleus, we then aimed to understand how the thalamocortical synaptic inputs associated with these transthalamic pathways compare with those of the corresponding corticocortical pathways (e.g. V1→pulvinar→LM vs. V1→LM inputs within LM cortex). Elucidation of the comparative synaptic impact of transthalamic and corticocortical pathways within the cortex first requires knowledge of how these pathways terminate across the laminar depth of their cortical target areas. Thus, we labeled corticocortical and transthalamic pathways originating from the same cortical source area with synaptophysin-mRuby to survey the distributions of their synaptic bouton terminations across the depth of their cortical target area. To label corticocortical pathways, injections of AAV-synaptophysin-mRuby were made in the cortical source area, while for transthalamic pathways, AAV1-Cre injections in the cortical source area were accompanied by AAV-FLEx-synaptophysin-mRuby in the pulvinar (Fig. 2A). We then analyzed the intensity of the mRuby labeling as a function of cortical depth within the appropriate target cortical region for each corticocortical and transthalamic pathway (see Materials and Methods) (Fig. 2B).

**Figure 2.**
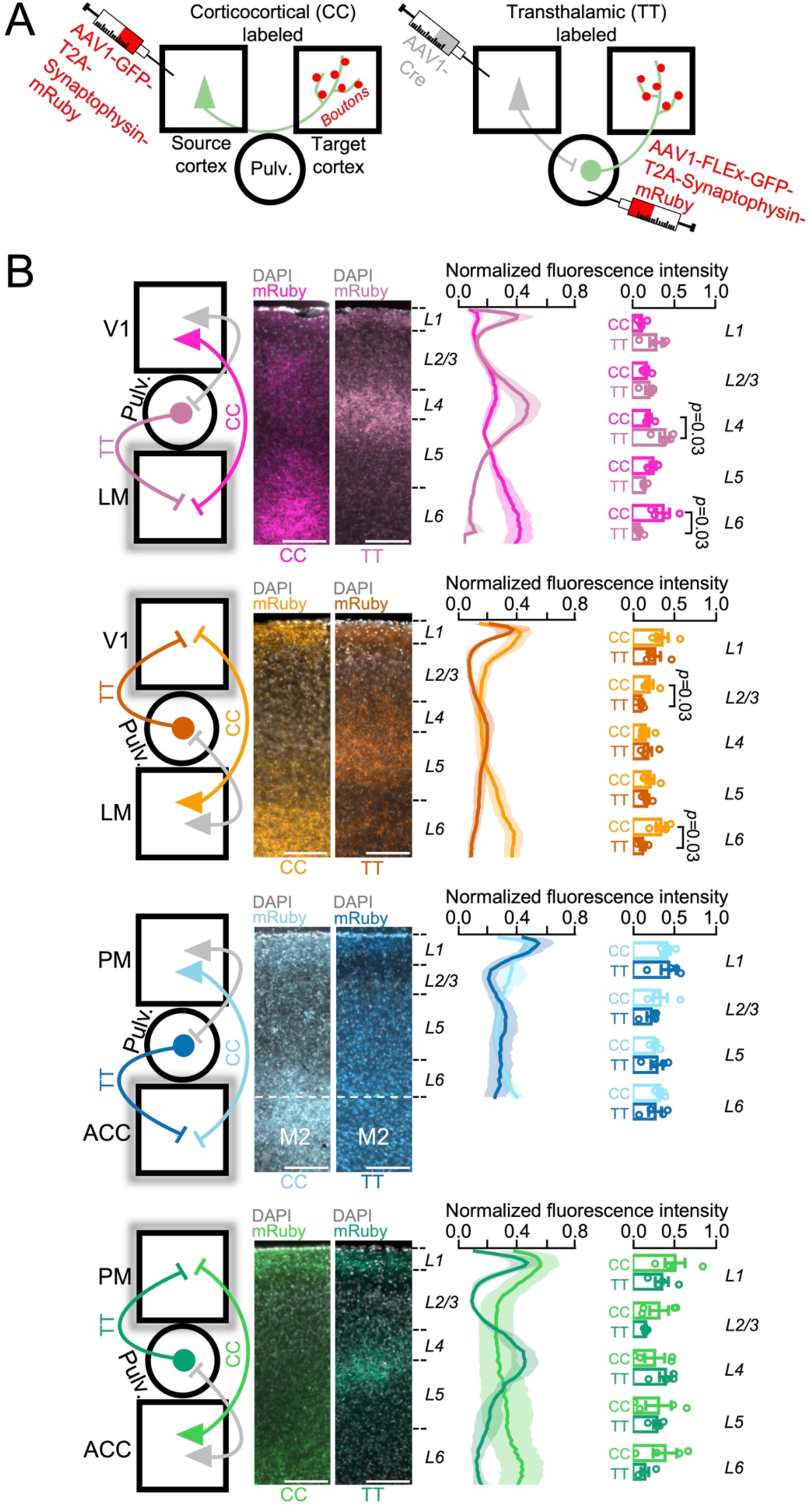
Laminar distributions of corticocortical and transthalamic synaptic terminations. (**A)** Schematic of viral vector injections for labeling either source-specific corticocortical (CC) or source-specific transthalamic (TT) synaptic boutons with synaptophysin-mRuby. **(B)** Quantification of the relative intensity of CC and TT synaptic boutons across the laminar depth of their postsynaptic cortical target region (*N=4* mice for each CC and TT pathway). Left plots are of normalized mRuby fluorescence intensity as a function of cortical depth for each pathway. Right plots compare normalized mRuby fluorescence intensity of CC and TT pathways in specific layers of the cortical target area (p-values from Mann-Whitney tests). For the pathways from PM to ACC, demarcation between ACC and secondary motor cortex (M2) is indicated in images. Scale bars = 150 µm.

For the initial feedforward pathway from V1 to LM, the relative densities of the synaptic terminations across the cortical depth of LM were largely distinct for the corticocortical and transthalamic projections. While the transthalamic V1→pulvinar→LM terminations most strongly clustered around layer 4 of LM, those of the corticortical V1→LM projections were found most abundantly in the infragranular layers, especially layer 6 (Fig. 2B). In the feedback direction (from LM to V1), the transthalamic LM→pulvinar→V1 and corticocortical LM→V1 projections also exhibited distinct laminar termination patterns, with the latter having a relatively higher termination density in both layer 2/3 and layer 6 (Fig. 2B). Contrastingly, the transthalamic and corticocortical projections in both feedforward and feedback directions between cortical areas PM and ACC exhibited comparable termination densities; no significant differences in relative terminal fluorescence intensity were found between the PM→pulvinar→ACC and PM→ACC projections or between the ACC→pulvinar→PM and ACC→PM projections (Fig. 2B).

### Dual-color optogenetic activation of corticocortical and transthalamic pathways reveals widespread monosynaptic convergence

Our analysis of the laminar distributions of synaptic bouton locations from the corticocortical and transthalamic pathways connecting both V1 with LM and PM with ACC indicate that, while a given corticocortical pathway often makes distinct, layer-specific synaptic terminations compared with its corresponding transthalamic counterpart, there are also ample anatomical opportunities for these pathways to converge within the same cortical layer, potentially monosynaptically on individual cortical neurons. Yet, the spatial colocalization of distinct synaptic terminations does not necessarily guarantee monosynaptic convergence; previous work has demonstrated cell-type-specific targeting by distinct presynaptic sources even when these sources co-terminate in a way that would physically allow for such convergence (Lafourcade *et al*., 2022; Perin *et al*., 2011; White, 2007; Song *et al*., 2005; Yoshimura *et al*., 2005). Thus, a fundamental question arises: Are corticocortical and transthalamic pathways integrated at the single-neuron level or do these pathways selectively target distinct cortical neurons, such that information from these pathways is instead integrated polysynaptically (Fig. 3A)? To answer this question, we expressed two spectrally distinct opsins (ChR2 and ChrimsonR) in corticocortical and transthalamic axons originating from the same cortical source (Fig. 3B), allowing us to optically activate these axons independently with 450-nm and 635-nm LED light in the same slice while performing whole-cell patch-clamp recordings of evoked excitatory postsynaptic currents (EPSCs) from cortical neurons. While ChR2 exhibits no activation by 635-nm light, ChrimsonR is activated by both 635- and 450-nm light (Klapoetke *et al*., 2014). However, by activating the ChrimsonR first with a long (250-ms) 635-nm light pulse, it will inactivate, rendering it unresponsive to a subsequent 450-nm light pulse that will only activate ChR2 (Hooks *et al*., 2015). Thus, 635-nm and 450-nm light pulses presented in sequence allow for selective optical activation of axon terminals expressing ChrimsonR and ChR2, respectively, which with TTX and 4-AP in the bath solution (Petreanu *et al*., 2009) tests for monosynaptic convergence of distinct presynaptic input sources onto a recorded neuron.

**Figure 3.**
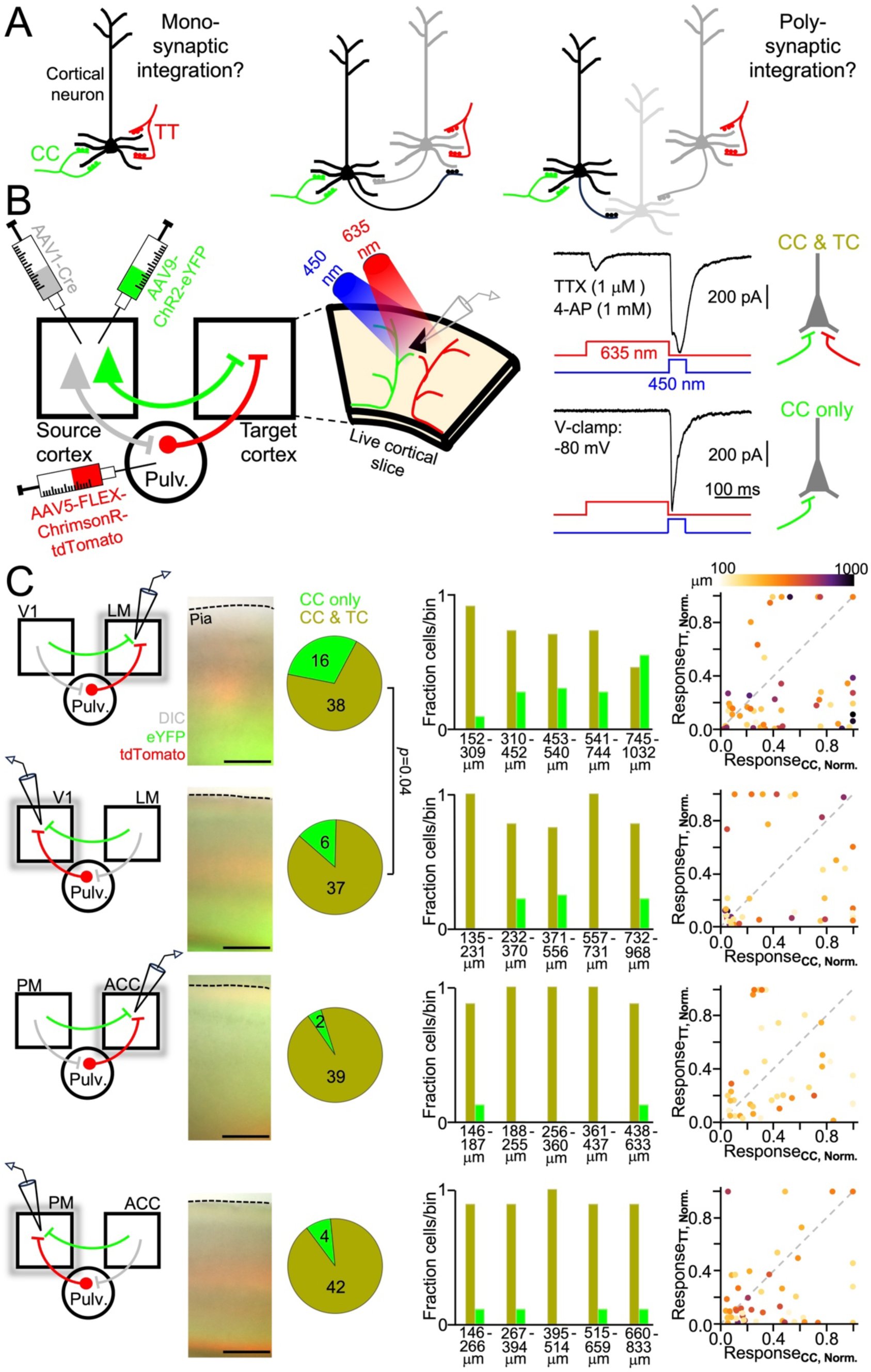
Dual-color optogenetic mapping of the monosynaptic convergence of corticocortical and transthalamic pathways. **(A)** Illustration of the possible circuit mechanisms of the integration of information from corticocortical (CC) and transthalamic (TT) pathways. **(B)** (*Left*) Schematic of viral vector injections and electrophysiological recordings combined with dual-color optogenetic axon terminal stimulation for testing monosynaptic convergence of CC and TT pathways originating from the same cortical source area. (*Right*) Examples of optically excitatory postsynaptic currents of a neuron receiving monosynaptic input from both pathways (top trace) or only one pathway (bottom trace). **(C)** Summary data of monosynaptic connectivity of CC and TT pathways, with data from each pathway grouping organized horizontally (pathways from V1 to LM: *N=7* mice, *n=54* neurons; pathways from LM to V1: *N=S* mice, *n=43* neurons; pathways from PM to ACC: *N* = 4 mice, *n* = 41 neurons; pathways from ACC to PM: *N=4* mice, *n=46* neurons). Images show examples of live cortical slices with both CC and TT axons labeled within them (scale bars= 300 µm). For each CC/TT pathway combination, pie charts indicate total counts of neurons receiving either both CC and TT input (yellow) or CC input alone (green). Cell count data are further broken into bins of laminar depth from the pia, with bin delimiters based on quintiles of the depth distributions for recorded neurons. Rightmost plots are of normalized excitatory charge recorded from optical stimulation of CC and TT axons for each recorded neuron, color-coded by its laminar depth.

Using dual-color optical stimulation of transthalamic and corticocortical axon terminals originating from the same cortical source region while recording postsynaptic responses from neurons in the cortical target region, we find that monosynaptic convergence of these two information streams is a prevalent feature at multiple points of the visual sensory processing hierarchy. Yet, the extent to which monosynaptic transthalamic-corticocortical convergence manifested within the cortex differed with hierarchical level. At the first stage of feedforward transmission from V1 to LM, ∼70% of all recorded LM neurons responded to optical activation of both V1→pulvinar→LM and V1→LM axon terminals (Fig. 3C). With the exception of LM neurons located in cortical layer 6, the majority of neurons throughout the laminar depth of LM exhibited monosynaptic convergence of the transthalamic and corticocortical pathways originating from V1 (Fig. 3C). In the feedback direction from LM to V1, monosynaptic convergence of transthalamic and corticocortical pathways was even more prevalent, with ∼86% of V1 neurons postsynaptically responsive to stimulation of each pathway, significantly higher than in the feedforward direction (*p =* 0.04, Fisher’s exact test) (Figure 3C). Furthermore, the majority of neurons in all layers of V1 displayed monosynaptic convergence of transthalamic and corticocortical pathways from LM (Figure 3C). For both feedforward and feedback transmission between PM and ACC, the overwhelming majority of cortical neurons responded monosynaptically to transthalamic and corticocortical axon stimulation (∼95% in the feedforward direction from PM to ACC and ∼91% in the feedback direction from ACC to PM) (Fig. 3C). This strong preponderance of pathway convergence featured in all cortical layers of PM and ACC (Fig. 3C).

We also tested whether the monosynaptic input magnitude of a transthalamic pathway corresponded with that of its associated corticocortical pathway (e.g. whether LM cortical neurons that exhibited large V1→pulvinar→LM responses also exhibited large V1→LM responses). For each recorded neuron in a cortical slice, we calculated the total excitatory charge passed during 635-nm LED stimulation and 450-nm LED stimulation and normalized these values to the maximum such charges recorded in the slice (see Materials and Methods). Thus, each recorded cortical neuron had a (Response_CC,Norm._, Response_TT,Norm._) data pair, which was plotted to determine whether transthalamic input strength was correlated with corticocortical input strength at the single-neuron level (Fig. 3C). Overall, we observed weak correlations between the monosynaptic input strengths of transthalamic and corticocortical pathways (Fig. 3C) (Spearman’s rank correlation coefficients [ρ] and *p-*values – V1-to-LM pathways: ρ = 0.30, *p* = 0.03; LM-to-V1 pathways: ρ = 0.21, *p* = 0.2; PM-to-ACC pathways: ρ = 0.27, *p* = 0.3; ACC-to-PM pathways: ρ = 0.14, *p* = 0.4). Thus, while transthalamic-corticocortical monosynaptic convergence is widespread across many hierarchically defined visual cortical regions, the input strength of one pathway does not generally predict that of the other pathway.

### Synaptic physiology of hierarchically defined transthalamic and corticocortical pathways

Having determined that a fundamental feature of source-specific transthalamic and corticocortical pathways is that they predominately engage in convergent monosynaptic activation of individual cortical neurons, we next sought to determine whether these pathways are physiologically comparable or distinct. While the synaptic physiology of first-order thalamic and intracortical synapses within primary sensory cortical areas is well-established (Kloc and Maffei, 2014; Cruikshank *et al*., 2010; Beierlein *et al*., 2003; Beierlein and Connors, 2002; Gil *et al*., 1999 & 1997), we lack commensurate knowledge for interareal synaptic connections within the cortex, particularly the characteristics of hierarchically defined corticocortical and transthalamic linkages.

To study the postsynaptic responses of cortical neurons to sustained activation of source-specific transthalamic and corticocortical pathways, we selectively labeled the axon terminals of these pathways with ChR2 using viral vector methods and prepared cortical slices to record the excitatory postsynaptic currents elicited by 10-Hz stimulation of the ChR2-expressing terminals with 450-nm LED light (Fig. 4A). We used voltage-clamp recordings at very hyperpolarized or very depolarized command potentials (along with the appropriate pharmacological blockers in the bath solution) to isolate either AMPA-based responses or NMDA-based responses, respectively (Fig. 4A), which we then converted to AMPA- or NMDA-based conductance responses (see Materials and Methods). For every cortical neuron recorded, the intensity of the 450-nm LED light was titrated to be 1.5x the threshold for eliciting an AMPA-based response, allowing us to compare data across different neurons.

**Figure 4.**
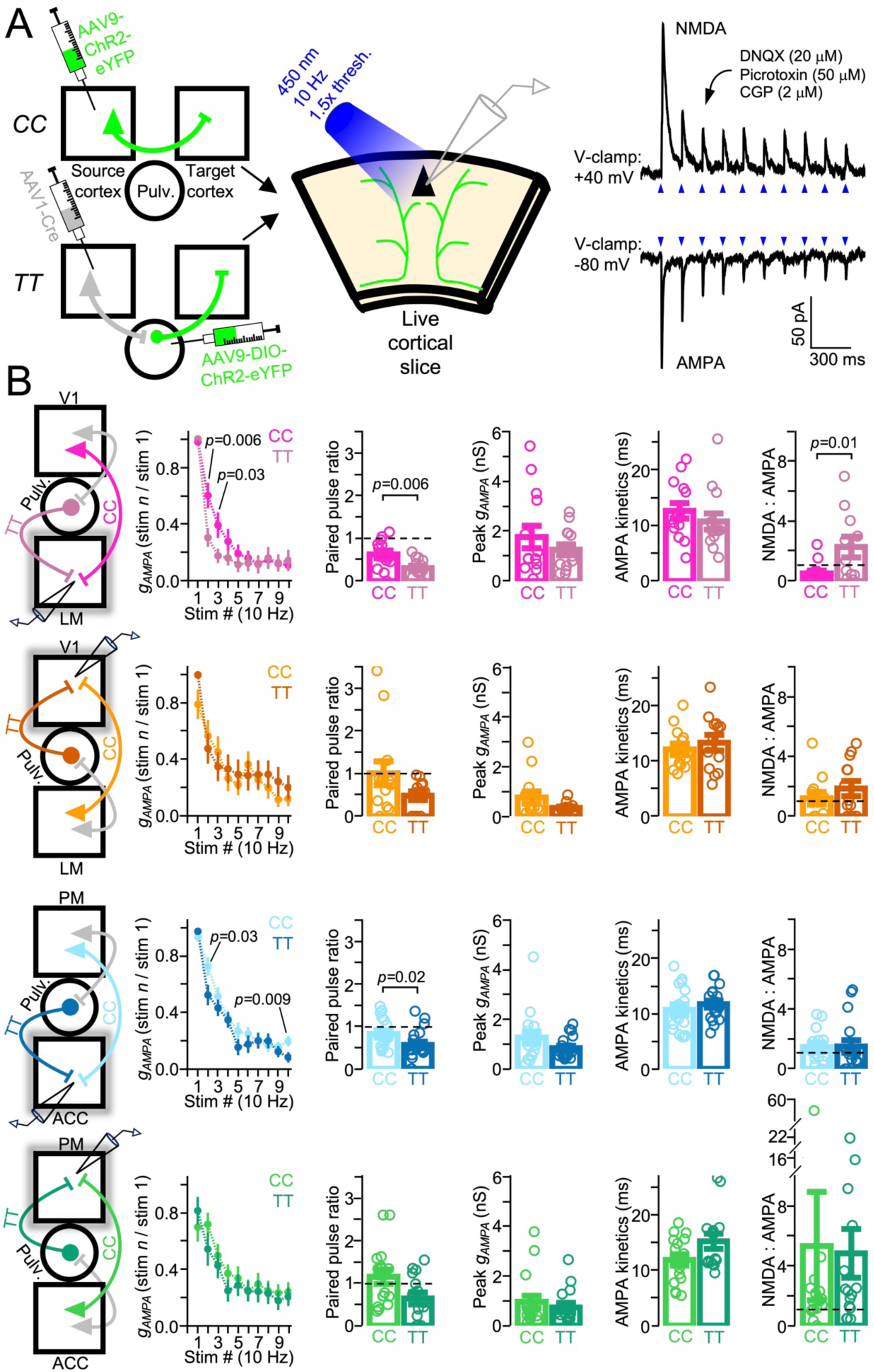
Synaptic physiology of corticocortical and transthalamic pathways. **(A)** (*Left*) Schematic of viral vector injections and electrophysiological recordings combined with repetitive axon terminal stimulation for studying the physiology of the glutamatergic synapses made by hierarchically distinct corticocortical (CC) and transthalamic (TT) pathways. (*Right*) Example of the AMPA and NMDA synaptic currents evoked in a recorded cortical neuron upon 10-Hz stimulation of ChR2-labeled axon terminals with 450-nm LED light. **(B)** Summary data of the synaptic properties compared between CC and TT pathways linking different cortical areas, with data from each CC/TT pathway combination organized horizontally. V1→LM: *N=7* mice, *n*=14 neurons; V1→pulv.→LM: *N*=*B* mice, *n*=14 neurons; LM→V1: *N=5* mice, *n*=13 neurons; LM→pulv.→V1: *N=5* mice, *n*=13 neurons; PM→ACC: *N=7* mice, *n*=17 neurons; PM→pulvinar→ACC: *N=6* mice, *n=20* neurons; ACC→PM: *N=5* mice, *n*=17 neurons; ACC→pulv.→PM: *N=5* mice, *n*=15 neurons. *P*-values from Mann-Whitney tests.

We aimed to clarify whether transthalamic and corticocortical pathways originating from the same cortical source region have similar or distinct functional impacts on neurons in a cortical target region. We thus compared several fundamental synaptic properties between the aforementioned pathways (i.e. V1→pulvinar→LM vs. V1→LM, LM→pulvinar→V1 vs. LM→V1, PM→pulvinar→ACC vs. PM→ACC, ACC→pulvinar→PM vs. ACC→PM): paired-pulse ratio, peak evoked AMPA conductance, evoked AMPA conductance kinetics, and the ratio of the evoked NMDA to evoked AMPA conductance. The laminar depths of the cortical neurons recorded during the stimulation of each pathway were comparable (V1→pulvinar→LM vs. V1→LM: 362 ± 14 μm vs. 329 ± 18 μm, *p* = 0.1; LM→pulvinar→V1 vs. LM→V1: 226 ± 12 μm vs. 231 ± 11 μm, *p =* 0.6; PM→pulvinar→ACC vs. PM→ACC: 201 ± 8 μm vs. 196 ± 10 μm, *p* = 0.9; ACC→pulvinar→PM vs. ACC→PM: 234 ± 7 μm vs. 214 ± 7 μm, *p* = 0.06; Mann-Whitney tests). In the initial feedforward processing stage from V1 to LM, we find that while transthalamic and corticocortical pathways activate LM neurons with similar strength and activation time-course (i.e. peak *g_AMPA_* and AMPA kinetics), the responses evoked by the transthalamic pathway were significantly more depressing and with a significantly higher NMDA component to the response (Fig. 4B). In the feedback direction from LM to V1, while transthalamic and corticocortical pathways elicited smaller AMPA-based postsynaptic responses (peak *g_AMPA_*) than in the feedforward direction (V1→pulvinar→LM vs. LM→pulvinar→V1: 1.3 ± 0.2 nS vs. 0.3 ± 0.06 nS, *p* = 0.0007; V1→LM vs. LM→V1: 1.8 ± 0.5 nS vs. 0.8 ± 0.2 nS, *p =* 0.02; Mann-Whitney tests), the LM→pulvinar→V1 and LM→V1 pathways were comparable in the synaptic dynamics, AMPA-based response magnitude and kinetics, and NMDA-to-AMPA ratio that they elicited in V1 neurons (Fig. 4B). For the feedforward connections between the higher-order cortical areas PM and ACC, activation of the transthalamic pathway elicited more synaptic depression in ACC neurons compared to the corticocortical pathway, but with otherwise comparable synaptic characteristics (Fig. 4B). Finally, the transthalamic and corticocortical pathways from ACC to PM were comparable in all synaptic characteristics analyzed (Fig. 4B).

## DISCUSSION

The synaptic pathways interconnecting distinct regions of the neocortex are critical for many higher-level brain operations. While direct corticocortical connections between cortical regions have traditionally been considered the principal mode for long-range cortical processing, indirect transthalamic pathways involving higher-order thalamus have increasingly been appreciated for their powerful influence over dynamic cortical functional connectivity. Yet, our knowledge of the comparative synaptic impacts of transthalamic and corticocortical pathways at distinct hierarchical processing stages is extremely limited, preventing a mechanistic understanding of how these two pathways jointly contribute to cortex-wide dynamics. Here, using the mouse visual system and the pulvinar nucleus as a model of hierarchical cortical sensory processing, we have uncovered some of the synaptic mechanisms by which specific corticocortical and transthalamic pathways functionally influence cortical neurons.

We first presented a strategy by which specific visual transthalamic pathways can be genetically targeted, labeling pulvinar neurons by both their corticothalamic input source and their thalamocortical output target. This novel technique revealed that while the thalamic neurons linking early visual cortical areas V1 and LM in the feedforward and feedback directions are spatially segregated within the pulvinar, those linking PM and ACC (two higher-order cortices) are largely intermingled in thalamic space. This result potentially indicates that transthalamic pathways at lower hierarchical processing levels are more direct relays of inputs from single cortical areas whereas those at higher hierarchical levels can sample and integrate inputs from multiple cortical sources. A substantial body of previous work in both primates (Cortes *et al*., 2026 & 2024) and rodents (Bennett *et al*., 2019; Zhou *et al*., 2017) has subdivided the pulvinar into multiple functionally and anatomically defined zones. Our method for labeling higher-order thalamic neurons based on the transthalamic pathway they serve offers an additional dimension of functional parcellation of not only the pulvinar, but other higher-order thalamic nuclei as well. Relatedly, recent work applying monosynaptic rabies tracing to distinct, cortical projection target-specific pulvinar neurons indicates that these neurons receive presynaptic inputs from a diverse collection of both cortical and subcortical regions (Leow *et al*., 2022; Blot *et al*., 2021). Yet, with target-specific labeling alone, it is not clear whether each labeled pulvinar neuron represents a distinct transthalamic pathway with a distinct cortical source or if each pulvinar neuron in fact integrates corticothalamic input from multiple sources. By labeling pulvinar neurons based not only on their thalamocortical projection target but also their corticothalamic projection source, as we have introduced here, future work will be able to distinguish between two possibilities regarding the fundamental synaptic logic of higher-order thalamic areas: whether transthalamic pathways are essentially channels for relaying information from one cortical region to another or whether higher-order thalamic neurons are integrative units that perform computations on signals originating from multiple cortical regions. This crucial question can be addressed by future work using either physiological or anatomical methods, such as by recording the postsynaptic responses in Flp-labeled pulvinar neurons (Fig. 1C) when optically stimulating distinct corticothalamic source axons or performing Flp-dependent monosynaptic rabies tracing (i.e. cTRIO [Schwarz *et al*., 2015]), respectively.

Using dual-color optogenetic circuit-mapping, we determined that a pervasive feature of interareal cortical synaptic communication within the mouse visual system is the monosynaptic integration of transthalamic and corticocortical pathways by individual cortical neurons. Since corticocortical and subcortically projecting neurons (e.g. corticothalamic neurons) are largely separate classes that also avoid synapsing with one another (Petrof *et al*., 2012; Brown and Hestrin, 2009), the cortical neurons in the target region that receive the higher-order thalamocortical and corticocortical inputs likely constitute the first location at which transthalamic and corticocortical information (e.g. V1→pulvinar→LM and V1→LM) is combined. Given that this information is largely integrated at the single neuron level, as described here, it will be important to determine the precise subcellular mechanisms by which transthalamic and corticocortical inputs are combined along the somatodendritic axis to influence the spiking output of the recipient cortical neurons. Given the complex relationship between the electrogenic properties of cortical neuron dendrites and the location of synaptic inputs (Harnett *et al*., 2012; London and Häusser, 2005; Reyes, 2001), it will be critical to determine the precise somatodendritic locations of transthalamic and corticocortical terminals on cortical neurons and develop experimental preparations to determine the resultant synaptic integration principles, such as whether integration is linear or supralinear and how the integration depends on the temporal delays between the inputs (Rindner *et al*., 2022). Studies of the mouse primary somatosensory cortex suggest that the role of feedback higher-order thalamocortical inputs is to gate the coupling between sensory-related signals arriving near the soma and basal dendrites and contextual signals arriving at the apical dendrites (Suzuki and Larkum, 2020), rather than integrating with cortical inputs *per se*. Future investigations of both feedforward and feedback transthalamic pathways will need to determine if higher-order thalamocortical inputs serve similar or distinct circuit functions at different hierarchical locations.

The combined properties of release probability, number of release sites from presynaptic terminals, and the number and type of receptors on the postsynaptic neuron establish the overall functional impact of synaptic pathways under different states of activation (e.g. low-vs. high-frequency action potentials in the axon terminal). The vast majority of our knowledge of activity-dependent synaptic efficacy of cortical and thalamic synapses comes from studies of first-order thalamic nuclei and primary sensory cortices. This previous work has converged on a consensus view that first-order thalamocortical synapses within primary sensory cortex are very strong with highly depressing synaptic dynamics, whereas individual intracortical synapses are relatively weak at low presynaptic activation frequencies, with steady or facilitating dynamics. These properties are thought to be suited to the distinct roles of these synapses within primary sensory cortex, with thalamocortical synapses providing a high-fidelity signal of the sensory content of the periphery and intracortical synapses further modifying this content to extract higher-level features (e.g. orientation and figure-ground segregation) or to modulate the gain of the signal in a state-dependent way (Niell and Scanziani, 2023; Ferguson and Cardin, 2020). Given the prevailing view that long-range transmission of signals across different cortical areas relies on direct corticocortical synapses, it would stand to reason that these synapses might have analogous properties to those of first-order thalamocortical synapses in primary sensory cortex. Yet, this view, as well as whether the accompanying transthalamic synapses share similar properties, is uncertain. Here, we showed that transthalamic synapses at the first intercortical transmission stage from V1 to LM exhibit more highly depressing synaptic dynamics compared to the direct corticocortical projections between those areas, whereas the feedback transthalamic and corticocortical inputs from LM to V1 were largely comparable and less depressing. These findings complement recent results reported in both the mouse visual and somatosensory cortex (Miller-Hansen and Sherman, 2022). More synaptic depression from the transthalamic pathway compared to the corticocortical pathway was also a feature of the feedforward connection between cortical areas PM and ACC. Therefore, a possible role of feedforward transthalamic pathways may be to “prime” the target cortical neurons for incoming corticocortical synaptic barrages. Interestingly, for all pathways we studied, transthalamic and corticocortical pathways originating from the same cortical source elicited AMPA-based responses of comparable magnitude and kinetics in the postsynaptic neurons of the target cortex. This result suggests that transthalamic and corticocortical pathways, while dynamically distinct in the feedforward direction, activate synapses with similar numbers or types of AMPA receptors with similar locations on the somatodendritic axis. Though, definitive physiological evidence of this would require more detailed quantal analyses of the responses and attenuation-based analyses for estimations of electrotonic distances of the synapses (Häusser and Roth, 1997). Another important distinction between the feedforward transthalamic and corticocortical pathways from V1 to LM, but not other pathways, was that activation of the transthalamic pathway elicited an excitatory response with a significantly stronger NMDA component. Given the prolonged kinetics and calcium-permeability of NMDA compared to AMPA conductances, transthalamic synapses from V1 to LM might offer a potent window of opportunity to permit the effective activation of LM neurons both acutely and potentially for long-term potentiation (LTP) of the pulvinar thalamocortical inputs. Indeed, higher-order thalamocortical inputs to sensory cortex have previously been shown to undergo LTP selectively during specific sensory experiences (Audette *et al*., 2019; Gambino *et al*., 2014).

It has been postulated that any cortical region receiving direct input from a given source cortical region also receives disynaptic input from the same source via higher-order thalamus (Shipp, 2003). The functional significance of these seemingly duplicate connections remains one of the most pressing questions regarding cortical and thalamic systems (Sherman and Guillery, 2013). One possibility is that transthalamic pathways carry distinctly processed sensory signals compared to those carried by corticocortical pathways, such that their integration gives rise to a coherent percept. Alternatively, transthalamic pathways might fundamentally engage in sensorimotor transformations by virtue of the branching axons of their corticocothalamic component (Guillery, 2003). Finally, rather than strictly engaging in sensory processing, transthalamic activity might serve as a “functional connectivity signal” to dynamically alter information flow within the cortex. Our results provide a much-needed synaptic basis for further exploration of these distinct functional roles.

## ACKNOWLEDGEMENTS

This study was supported by NIH grant R00 EY-030550.

## MATERIALS AND METHODS

### Animal details

For electrophysiological recordings and synaptic bouton labeling analyses, C57BL/6J mice of both sexes (The Jackson Laboratory, Strain #: 000664) aged 7–12 weeks were used. For localization of transthalamic relay cells within the pulvinar nucleus, heterozygous Ai65f mice of both sexes (The Jackson Laboratory, B6.Cg-Gt(ROSA)26Sor^tm65.2(CAG-tdTomato)Hze^/J, Strain #: 032864) aged 7-12 weeks were used. For some experiments used for testing the transsynaptic infection properties of AAV1-Cre (see *Viruses* and Fig. 1B), we used heterozygous Ai9 mice (The Jackson Laboratory, B6.Cg-Gt(ROSA)26Sor^tm9(CAG-tdTomato)Hze^/J, Strain #: 007909). All procedures involving mice were approved by the University at Buffalo Institutional Animal Care and Use Committee.

### Surgical procedures

For all experiments, transgenes were introduced into neurons of interest via intracranial virus injection into the cortex and thalamus of anesthetized mice. Mice were anesthetized on a stereotaxic apparatus with 0.8 –1.5% isoflurane in O_2_. Burr holes were drilled into the cranium at the appropriate stereotaxic coordinates for the various cortical and thalamic regions of interest, as measured by a digital micropositioning system (Kopf Instruments Model 940). Pulled glass micropipettes were filled with virus solution and lowered into the brain at the desired depth relative to the pia, with virus then injected into the brain at 3–7 nL/min using a microinjection system (Stoelting Quintessential Stereotaxic Injector). Micropipettes were left in place for ∼5 min before withdrawing.

### Stereotaxic coordinates for virus injections

This study concerns the synaptic interconnectivity of four mouse cortical regions (primary visual cortex [V1], lateromedial visual cortex [LM], posteromedial visual cortex [PM], and anterior cingulate cortex [ACC]) with the higher-order mouse visual thalamus (the lateral posterior nucleus of the thalamus [LP], or pulvinar). For V1 injections, 2 injection sites were used with different volumes injected: (from bregma, in mm) -2.5 (*x*), -3.5 (*y*), -0.6 (*z*) [200 nL] and -2.5 (*x*), -2.8 (*y*), -0.6 (*z*) [50 nL]. For LM injections, 3 injection sites were used with injection volumes of 100 nL each: (from bregma, in mm) -3.8 (*x*), -4.3 (*y*), -0.6 (*z*); -3.8 (*x*), -3.8 (*y*), -0.6 (*z*); and -3.8 (*x*), -3.2 (*y*), -0.6 (*z*). For PM injections, 2 injection sites were used with different volumes injected: (from bregma, in mm) -1.8 (*x*), -2.7 (*y*), -0.6 (*z*) [50 nL] and -1.5 (*x*), -2.9 (*y*), -0.6 (*z*) [100 nL]. For ACC injections, 4 injection sites were used with injection volumes of 100 nL each: (from bregma, in mm) -0.3 (*x*), +0.14 (*y*), -0.9 (*z*); -0.3 (*x*), -0.1 (*y*), -0.9 (*z*); -0.3 (*x*), -0.6 (*y*), -0.9 (*z*); -0.3 (*x*), -1.1 (*y*), -0.9 (*z*). For pulvinar injections, injection volumes were 200 nL and injection coordinates depended on the transthalamic pathway targeted: for the pulvinar-to-LM and pulvinar-to-V1 pathways (from bregma, in mm) -1.64 (*x*), -2.28 (*y*), -2.7 (*z*); for the pulvinar-to-ACC pathway (from bregma, in mm) -1.0 (*x*), -2.03 (*y*), -2.7 (*z*); for the pulvinar-to-PM pathway (from bregma, in mm) -1.6 (*x*), -2.06 (*y*), -2.7 (*z*).

### Viruses

All viruses were allowed at least 3 weeks to express before experimentation. For anterograde transsynaptic labeling of pulvinar neurons with Cre recombinase based on their corticothalamic source region, injection of high-titer (1×10^13^ GC/mL) AAV1-Syn-Cre (SignaGen, SL101461) was made within cortex (Zhou *et al*., 2022; Zingg *et al*. 2020 & 2017). Retrograde labeling of pulvinar neurons with thalamocortical terminals in specific cortical target regions was accomplished with the CAV2 virus, specifically CAV2-FLEx^loxP^-Flp (5×10^12^ pp/mL) (CRNS Plateforme de Vectorologie de Montpellier, CAV FlxFlp). For labeling of corticocortical or transthalamic synaptic boutons within a postsynaptic cortical target region, cortical injections of AAV1-hSyn-mGFP-T2A-Synaptophysin-mRuby (2×10^12^ GC/mL) (Biohippo, AAVX-1243-21-12) or thalamic injections of AAV1-hSyn-FLEx-mGFP-2A-Synaptophysin-mRuby (2×10^12^ GC/mL) (Addgene, 71760-AAV1) (accompanied by the appropriate cortical AAV1-Syn-Cre injections) were made, respectively. For dual-color optogenetic synaptic pathway mapping experiments in cortical slices, a 1:1 mixture of AAV1-Syn-Cre and AAV9-hSyn-hChR2-eYFP (2×10^12^ GC/mL) (Addgene, 26973-AAV9) was injected into the presynaptic cortical source region and AAV5-Syn-FLEX-rc[ChrimsonR-tdTomato] (2×10^12^ GC/mL) (Addgene, 62723-AAV5) was injected into the pulvinar. For analysis of the synaptic physiology of distinct corticocortical and transthalamic pathways, cortical injections of and AAV9-hSyn-hChR2-eYFP (2×10^12^ GC/mL) or thalamic injections of AAV9-EF1a-DIO-hChR2-eYFP (2×10^12^ GC/mL) (Addgene, 35509-AAV9) (accompanied by the appropriate cortical AAV1-Syn-Cre injections) were made, respectively. AAV9 was chosen as the serotype for introducing ChR2 into projection axons for synaptic physiology experiments based on the documented optimal preservation of native short-term synaptic dynamics when using this serotype combined with optogenetic stimulation of axon terminals in *ex vivo* slices (Martinetti *et al*., 2022; Jackman *et al*., 2014).

### Histology

Mice with viral injections for histological analysis were deeply anesthetized with isoflurane and transcardially perfused with PBS followed by 4% PFA (in PBS). Perfused brains were removed from the cranium and postfixed for 24 hr at 4°C. Coronal sections (50 μm) were then cut using a vibratome (Compresstome VF-510-OZ, Precisionary), after which free-floating sections were mounted to microscope slides, covered with mounting medium (ProLong Gold Antifade Mountant with DAPI, Invitrogen), and sealed with cover glass. Photomicrographs of mounted sections were taken with a Leica DFC9000 GT sCMOS camera mounted to an epifluorescence microscope (Leica DM 6B).

### Ex vivo slice preparation

Live brain slices (300 μm thick) were prepared from virally injected mice. Mice were first deeply anesthetized with isoflurane and transcardially perfused with an ice-cold NMDG-based cutting solution saturated with a 95% O_2_ / 5 % CO_2_ mixture. The composition of the solution was as follows (in mM): 93 NMDG, 2.5 KCl, 1.2 NaH_2_PO_4_, 30 NaHCO_3_, 20 HEPES, 25 dextrose, 5 Na-ascorbate, 2 thiourea, 3 Na-pyruvate, 0.5 CaCl_2_, 10 MgSO_4_. Following perfusion, brains were extracted and sectioned in this same solution with a vibratome (Leica VT 1000S). Brain slices were then transferred to an incubation chamber (maintained at ∼33°C) containing the NMDG-based solution. The concentration of Na^+^ was gradually raised within the incubation chamber over the first 20 min of incubation by adding increasing amounts of 1 M NaCl dissolved NMDG-based solution (0 min: [Na^+^]=45 mM, 5 min: [Na^+^]=52 mM, 10 min: [Na^+^]=65 mM, 15 min: [Na^+^]=87 mM, 20 min: [Na^+^]=129 mM) (Ting *et al*., 2018). After 30 min incubation in the modified NMDG-based solution, slices were transferred to an incubation chamber at room temperature containing artificial cerebrospinal fluid (ACSF) with the following composition (in mM): 126 NaCl, 3 KCl, 1.25 NaH_2_PO_4_, 26 NaHCO_3_, 10 dextrose, 2 CaCl_2_, 2 MgSO_4_, saturated with a 95% O_2_ / 5 % CO_2_ mixture. Slices remained in this incubation chamber until recording.

### Electrophysiological recordings

After at least 30 additional minutes in the final incubation chamber, slices were transferred to a recording chamber bathed continuously (∼5 mL/min) with ACSF maintained at 33°C. Cells were visualized with an Olympus BX51WI microscope equipped with IR-DIC and epifluorescence optics. Cortical pyramidal neurons were specifically targeted for this study based on their somatic taper toward the pia under IR-DIC visualization. Whole-cell patch-clamp recordings were performed on these cells using pulled borosilicate glass pipettes (tip resistances ∼5 MΩ) filled with a Cs-based internal solution containing the following (in mM): 130 Cs-gluconate, 4 CsCl, 2 NaCl, 10 HEPES, 0.2 EGTA, 4 ATP-Mg, 0.3 GTP-Na, 14 phosphocreatine-2Na, 5 QX-314 (pH ∼7.25, ∼300 mOsm). Reported voltages were not corrected for the liquid junction potential. Whole-cell voltage-clamp recording data were filtered (DC–3 kHz) and digitized at 20 kHz using a MultiClamp 700B patch-clamp amplifier (Axon) and Cambridge Electronic Design (CED) hardware and software. Series resistances (<30 MΩ) were continuously monitored and compensated offline. AMPA-based synaptic currents were recorded at a command potential of -80 mV and NMDA-based synaptic currents were recorded at a command potential of +40 mV with the following (in μM) bath-applied in the ACSF to block AMPA-, GABA_A_-, and GABA_B_-based responses at this potential: 20 DNQX, 50 picrotoxin, 2 CGP-55845, respectively. For recordings of NMDA-based currents, we first ensured that bath-application of the drug mixture abolished all AMPA-based currents at -80 mV at full intensity of the 450-nm LED. For dual-color optogenetic circuit mapping experiments, recordings were made in ACSF containing 1 μM TTX and 1 mM 4-AP to ensure that optically evoked responses were monosynaptic in nature (Petreanu *et al*., 2009).

### Optogenetic stimulation

All slices were first screened for the presence of opsin-expressing axon terminals under low-magnification epifluorescence. Collimated LED light for optical excitation (pE-400max, CoolLED) was reflected through a dichroic mirror (FF655-Di01, Semrock) and an immersion objective (LUMPLFLN40XW, Olympus), resulting in a spot diameter of ∼1 mm. A 450-nm LED was used to excite ChR2 and a 635-nm LED was used to excite ChrimsonR. For dual-color optogenetic circuit mapping experiments, maximum-intensity (36 mW/mm^2^) 635-nm LED light was applied for 250 ms to activate ChrimsonR, followed immediately by maximum-intensity (79 mW/mm^2^) 450-nm LED for 50 ms to activate ChR2 following the inactivation of Chrimson (Hooks *et al*., 2015) (see “Results”). For repetitive stimulation of ChR2-expressing axon terminals for synaptic physiology assessments, 1-ms pulses of 450-nm LED light were delivered at 10 Hz with an intensity of 1.5× threshold for evoking AMPA-based responses (2–21 mW/mm^2^).

### Experimental design and statistical analysis

“*N*” refers to number of animals and “*n*” refers to number of neurons. Unless otherwise noted, data are summarized as mean ± standard error of the mean, which is computed over the number of neurons. Nonparametric statistics were used for statistical comparisons between different neuron or synapse types in this study: Mann-Whitney or Wilcoxon’s matched pairs test for comparisons of two groups; Kruskal-Wallis or Friedman’s test for comparison of multiple groups, followed by Dunn’s post-hoc test and adjustment of *p-*values by the Benjamini-Hochberg procedure for control of the false discovery rate; and Fisher’s exact test for group comparisons of binary synaptic connectivity results. *p-*values < 0.05 were considered statistically significant.

### Data analysis: Cell counts of transthalamically labeled pulvinar neurons

Pulvinar neurons labeled with tdTomato in Ai65f mice injected with AAV1-Syn-Cre in a presynaptic source cortical region and CAV2-FLEx^loxP^-Flp in a postsynaptic target cortical region were identified manually in each histological section. Each tdTomato-labeled pulvinar neuron was assigned an *x-*, *y-*, and *z-*value (in mm) in stereotaxic coordinates. The *x-*value was the medial-lateral distance from the midline, the *y-*value was the anterior-posterior distance from bregma (using the section at which the hippocampus disappeared from view as a fiducial landmark corresponding to 1 mm posterior to bregma [Paxinos and Franklin, 2013]), and the *z-*value was the dorsal-ventral distance from a line tangential to the pial surface. Each pulvinar neuron from the four different transthalamic pathways targeted in this study were plotted in 3D coordinates and the location distributions in each anatomical axis were compared among the four pathways.

### Data analysis: Cortical laminar distributions of corticocortical and transthalamic synaptic boutons

Fluorescence intensity of synaptophysin-mRuby was measured in 10-μm bins aligned to the pial surface, extending to the white matter below cortical layer 6. Measured fluorescence intensities in each laminar depth bin per section were then subtracted by the autofluorescence intensity in a cortical region without mRuby expression. These resulting fluorescence intensities were then normalized to the maximum intensity measured across sections per each animal. For analysis of normalized mRuby fluorescence in specific cortical layers, numerical values of layer-specific depth demarcations for V1/LM/PM and ACC were taken from D’Souza *et al*. (2022) and Vogt and Paxinos (2014), respectively.

### Data analysis: Optically evoked synaptic currents and conductances from whole-cell recordings

Recordings of whole-cell current at each command potential in voltage-clamp were aligned to optical stimulation onsets. The recorded current (*I_recorded_*) was then subtracted by the mean current 100 ms before optical stimulation onset, yielding Δ*I_recorded_*. Recorded voltages (*V_recorded_*) were corrected for series resistance (*R_series_*) using the following equation to account for the time-dependent voltage drop across the series resistance,

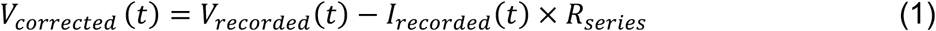

Corrected voltages were then similarly subtracted by the mean corrected voltage 100 ms before optical stimulation onset, yielding Δ*V_corrected_*. Optically evoked AMPA-mediated currents were calculated as follows,

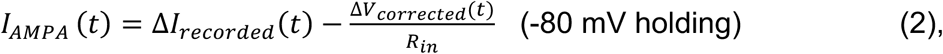

where *R_in_* is the input resistance of the cell (calculated from the cell’s voltage response to 5–10 pA current pulses in current-clamp), and is included for subtracting any current from Δ*I_recorded_* resulting from moving the cell away from its resting potential, thus isolating the synaptic current. After verifying that bath-application of DNQX, picrotoxin, and CGP-55845 fully abolished synaptic currents at -80 mV holding potential, optically evoked NMDA-mediated currents were similarly calculated,

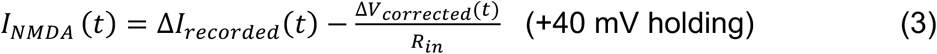

To compute AMPA and NMDA synaptic conductances (*g_AMPA_* and *g_NMDA_*) from recorded synaptic currents, the following equations were used,

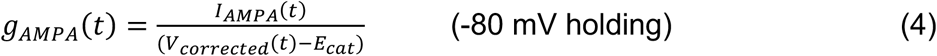

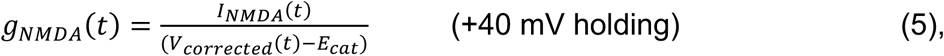

where *E_cat_* is the reversal potential for AMPA/NMDA cationic current previously measured using our Cs-based internal solution (*E_cat_* = +7 mV) (Neske *et al*., 2015). For each recorded cell, 10 1-ms, 10-Hz, 450-nm LED light pulses (see “Optogenetic stimulation”) were given every 15 s (∼10 trials), and the optically evoked AMPA- and NMDA-mediated responses were then trial-averaged. The peak AMPA or NMDA conductance was taken as the maximum of the mean *g_AMPA_* or *g_NMDA_* calculated during the 10-Hz optical stimulus trains. To characterize synaptic dynamics, the peak of the mean conductance was determined within a 10-ms window following each of the 10 optical stimulus pulses during the 10-Hz train. Each conductance value per pulse was then normalized to the maximum conductance value measured across all 10 pulses. Relatedly, the paired-pulse ratio was measured as the ratio of the peak conductance measured after the second pulse to the peak conductance measured after the first pulse. For each cell, the NMDA:AMPA ratio was calculated as the peak *g_NMDA_* measured during the 10-Hz train divided by the peak *g*_AMPA_ measured during the 10-Hz train. For quantification of conductance kinetics (a measure of the width of the excitatory postsynaptic conductance waveform), the following equation was used,

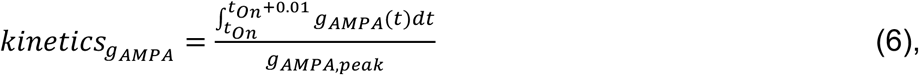

i.e., the area under the conductance waveform from *t_On_* to *t_On_* + 10 ms (where *t_On_* is the onset time of the optical pulse eliciting the largest conductance in the 10-Hz train) divided by the peak conductance value during that time period.

### Data analysis: Dual-color optogenetic circuit mapping of corticocortical-transthalamic pathway convergence

For experiments in which corticocortical and transthalamic pathways originating from the same cortical source were labeled with ChR2-eYFP and ChrimsonR-tdTomato, respectively, all slices were first screened to ensure that both eYFP- and tdTomato-expressing axons were present, ensuring the anatomical possibility of detecting monosynaptic convergence of the two presynaptic sources in candidate postsynaptic neurons. Whole-cell patch-clamp recordings (-80 mV command potential) were performed to record synaptic currents evoked by 635- and 450-nm light (max intensity) (see “Optogenetic stimulation”). To sample for connectivity across the entire cortical laminar depth, ∼6 cells per slice were recorded, spanning from just below layer 1 to the deepest part of layer 6. In order to recover the ChrimsonR in the transthalamic axons from inactivation for each trial, the intertrial interval between 635- and 435-nm light pulse sequences was 1 min (∼5 trials per cell). A cell was considered to have a monosynaptic connection from the transthalamic or corticocortical projection if the trial-averaged recorded current during the 635-nm light stimulus or 450-nm light stimulus, respectively, was significantly different from the trial-averaged recorded current 150 ms prior to the onset of the optical stimuli (Mann-Whitney test). To quantify the magnitudes of the transthalamic and corticocortical synaptic responses elicited during these dual-color circuit mapping experiments, the area under the trial-averaged recorded currents (i.e. synaptic charge, *Q_TT_* and *Q_CC_*) was computed during the 50 ms after 635- and 450-nm light stimulation, respectively. All transthalamic responses (*Q_TT_*) were normalized to the maximal transthalamic response recorded across all the cells from a given animal to yield a *Response_TT,Norm_* value per cell for analysis of the population dataset. Similarly, all corticocortical responses (*Q_CC_*) were normalized to the maximal corticocortical response recorded across all the cells from a given animal to yield *Response_CC,Norm_* for each cell.

